# Fluorosphere-Assisted Nano-Biosensor for Detection of Circulating microRNAs in Non-Small Cell Lung Cancer

**DOI:** 10.64898/2026.09.28.754875

**Authors:** Pradyumna Kumar Mishra, Ruchita Shandilya, Arpit Bhargava, Pooja Ratre, Vikas Gurjar, Olga Goryacheva, Rajnarayan Tiwari, Irina Yu Goryacheva

**Affiliations:** Division of Environmental Biotechnology, Genetics & Molecular Biology (EBGMB), ICMR-National Institute for Research in Environmental Health (NIREH), Bhopal, India; Faculty of Medical Research, Academy of Scientific and Innovative Research (AcSIR), Ghaziabad, India; Faculty of Science, Ram Krishna Dharmarth Foundation (RKDF) University, Bhopal, India; Institute of Chemistry, Saratov State University, Astrakhanskaya 83, 410012, Saratov, Russia; Faculty of Materials Science, Shenzhen MSU-BIT University, Shenzhen 518172, People’s Republic of China

**Keywords:** Circulating cell-free microRNA, Lung Cancer, NSCLC, Nano-hybrid array, Fluorosphere, Locked nucleic acid probe

## Abstract

Non-small-cell lung cancer (NSCLC) diagnosis is challenged owing to the need for reliable, non-invasive biomarkers. Circulating cell-free microRNAs (ccf-miRNAs) are promising candidates, but their low levels and short sequences in blood complicate detection. In this study, a novel nano-hybrid fluorescent array was developed to enable rapid and sensitive identification of NSCLC-related ccf-miRNAs. The array consists of poly(T)-modified fluorescent nano-polystyrene beads for capturing target miRNAs, and fluorophore-labeled, sequence-specific locked nucleic acid (LNA) probes for their detection. This design forms a sandwich structure with the target miRNA. Nano cytometry and fluorescence microscopy assessed both the capture and specificity of detection. The assay was tested on plasma samples for miR-16-5p and U6, evaluating selectivity, sensitivity, and reproducibility. The results showed that the poly(T)-modified beads efficiently captured the target miRNAs, and the LNA probes accurately distinguished the sequences. The assay enabled direct detection of miR-16-5p and U6 from plasma without amplification, demonstrating high selectivity, sensitivity, and reproducibility. Combining enrichment with nano-polystyrene beads and sequence-specific LNA probes addressed the main challenges of ccf-miRNA detection, namely their low abundance and short sequence length. As ccf-miRNAs are linked to NSCLC progression, this method could assist in early diagnosis, disease monitoring, and assessment of treatment response. The target-independent nature of the poly(T) capture layer allows for easy calibration to new miRNAs by changing the LNA probes, rendering it suitable for multiplex detection in clinical settings. However, further preclinical and clinical studies are needed before adoption in routine practice. This nano-hybrid fluorescent array presents a rapid and reliable approach for detecting low-abundance ccf-miRNAs in plasma. It’s encouraging performance and flexible design show potential for future application in NSCLC diagnosis and monitoring, subject to additional confirmation.

## 1 Introduction

Lung cancer (LC) remains one of the most frequently diagnosed malignancies worldwide, accounting for approximately 11.6% of all cancer cases and representing the leading cause of cancer-related mortality, contributing to nearly 18.4% of total cancer deaths globally (1, 2). Non-small-cell lung cancer (NSCLC) is the most prevalent subtype of lung cancer, accounting for approximately 85% of all diagnosed cases, and remains one of the leading causes of cancer-related mortality worldwide. The high incidence and mortality associated with NSCLC are largely attributed to delayed diagnosis, limited availability of reliable diagnostic tools, and the absence of well-organised screening programs (3). Although low-dose computed tomography (LDCT) has significantly improved early detection rates, several technical and clinical limitations, including high false-positive rates, radiation exposure, and cost-related concerns, restrict its widespread implementation as a routine screening strategy (4). Furthermore, conventional tissue biopsy procedures are invasive and often complicated by inadequate tissue acquisition, tumour heterogeneity, and procedural risks, thereby emphasising the need for minimally invasive and highly sensitive diagnostic approaches (5). During cellular turnover, apoptosis, necrosis, and active secretion processes, billions of cells release nucleic acids and other molecular constituents into the bloodstream. These circulating molecules serve as surrogate biomarkers that reflect the molecular landscape of the disease and can provide real-time information on tumour burden and treatment response (6).

Among the various circulating biomarkers, circulating cell-free microRNAs (ccf-miRNAs) have emerged as particularly promising candidates for cancer diagnosis and prognosis. ccf-miRNAs function as master regulators of gene expression and influence multiple tumorigenic pathways. Alterations in their expression levels often represent early molecular events preceding clinical manifestations of disease. Consequently, monitoring ccf-miRNA signatures may facilitate early cancer detection and provide insights into disease progression and treatment efficacy. These characteristics have established ccf-miRNAs as attractive molecular targets for liquid biopsy-based diagnostic applications (7, 8).

Despite their significant clinical potential, accurate detection and quantification of ccf-miRNAs remain challenging. Conventional analytical methods, including quantitative real-time PCR, microarrays, and next-generation sequencing, often require extensive sample processing, sophisticated instrumentation, complex probe designs, amplification steps, and considerable analysis time. Furthermore, issues related to reproducibility, sensitivity, and standardisation continue to limit their routine clinical implementation. Therefore, there is a growing need for simplified, rapid, and highly sensitive analytical platforms capable of detecting low-abundance circulating miRNAs with improved accuracy and reproducibility (9). Among the various signal transduction systems, nanomaterial-based platforms have attracted considerable attention owing to their unique physicochemical properties. Engineered nanomaterials exhibit high surface-to-volume ratios, enhanced optical properties, excellent stability, and versatile surface functionalization, making them ideal candidates for biosensing applications (10-12).

In particular, polymeric bead-based suspension arrays have emerged as powerful analytical tools for multiplexed biomarker detection. These systems enable simultaneous quantification of multiple analytes from small sample volumes and offer advantages such as high throughput, sensitivity, and ease of operation. Microsphere-based flow cytometry represents another promising approach for nucleic acid detection. Functionalized microspheres possessing carboxyl or amine groups can be chemically modified to immobilise oligonucleotides, proteins, peptides, and other biomolecules through covalent or non-covalent interactions (13). However, conventional carboxyl-modified latex microspheres often suffer from limitations, including particle aggregation, polymer degradation, fluorescence interference, poor microscopic visualisation, reduced reproducibility, and challenges associated with colocalization studies. These drawbacks can compromise analytical performance and hinder accurate biomarker quantification.

The present study describes a method for quantitative assessment of ccf-miRNAs in the isolated samples using a nano-polystyrene-based fluorescent array. Specifically, an oligonucleotide-nanopolystyrene-fluorophore-labelled locked nucleic acid-based flow cytometry method. The nano-hybrid assembly discussed herein provides a sensitive, direct method for quantifying ccf-miRNAs p by flow cytometry without amplification. The methodology requires the appropriate positioning of poly-T oligonucleotide sequences on the nano-polystyrene to effectively capture polyadenylated ccf-miR. The formed complex is then allowed to bind with customised fluorophore-labelled LNA (fLNA) probes to identify the specific ccf-miR. The fLNA probes are conformationally restricted nucleotide analogs that comprise of a ribose molecule in which 2′-oxygen and the 4′-carbon are associated with a methylene group to mimic RNA sugar conformation. The fLNA probes bind with the target miRNA molecules in a sequence-specific manner, abiding by the Watson−Crick base pairing rule, and exhibit high affinity towards the complementary target miRNA sequences. Such probes exhibit a significant melting temperature (Tm) difference between a perfectly matched ccf-miR of interest and a mismatched ccf-miR, thereby enabling good mismatch discrimination under optimal hybridisation conditions. The developed nano-hybrid assembly-based methodology enables rapid, reproducible, and real-time quantification of disease-specific ccf-miRs with high specificity, without the need for any complex & time-consuming amplification steps. The developed methodology might assist in highly sensitive and selective characterisation of the aberrantly expressed ccf-miRs in isolated samples, which would assist in the prediction of the altered biological processes for further analysis of different pathologies.

## 2. Materials and Methods

### 2.1. Reagents and Equipment

All the reagents used were of the highest available grade. Nucleospin® miRNA plasma kit for ccf-miRs isolation was procured from Macherey-Nagel, Germany. The poly-A-tailing kit was procured from Invitrogen (Thermo Fisher Scientific, US). PrimeScript 1st strand cDNA synthesis kit was obtained from Takara (Japan). Taq 2X mastermix was obtained from NEB (USA). CML latex-coated beads, EDC (1-ethyl-3-[3-dimethylaminopropyl] carbodiimide hydrochloride) and NHS (N-hydroxysuccinimide) were procured from Thermofisher Scientific (USA). MES (2-(N-morpholino) ethanesulfonic acid) was purchased from Sigma-Aldrich (USA). Tween 20 was procured from Bio-Rad Laboratories (USA). Allegra™ X-22R centrifuge was purchased from Beckman Coulter™. A Nanodrop 2000 spectrophotometer was purchased from Thermo Scientific. The GeneAmp® PCR system 9700 from Applied Biosystems was also utilised. The Attune NXT Flow cytometer was purchased from Thermo Fisher Scientific. A multimode reader (Dual double monochromator) was procured from Tecan Spark. The oligonucleotides listed in the table below were appropriately customised to meet the specified requirements **(Table 1)**.

**Table 1:**
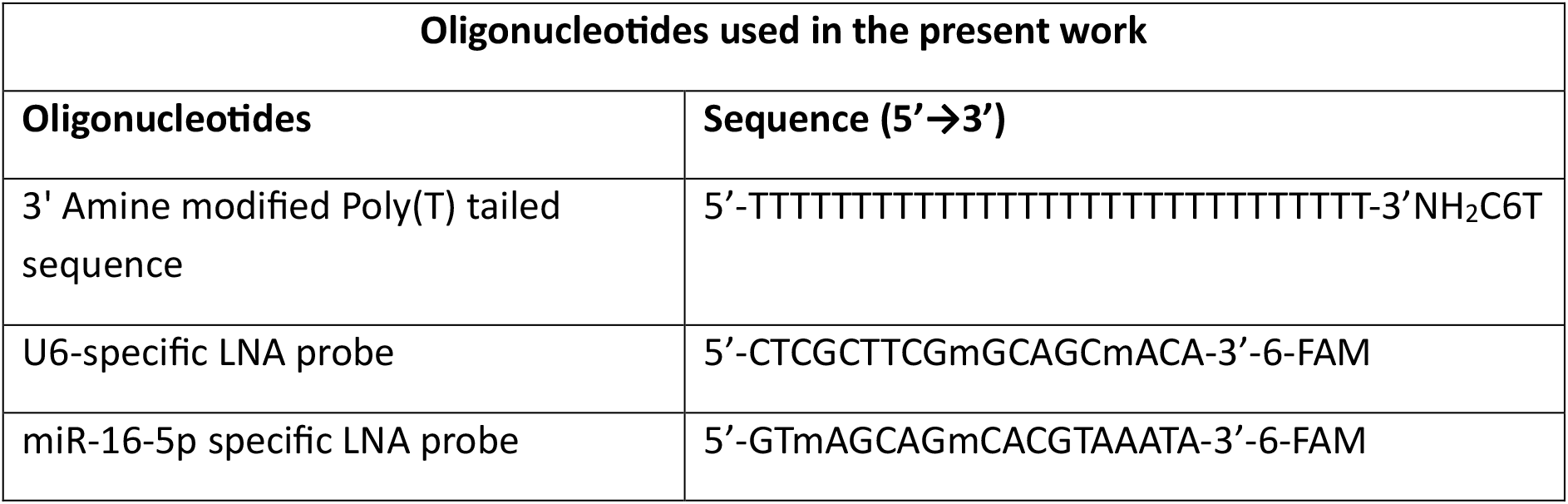
The customised oligonucleotides used for the study.

| Oligonucleotides used in the present work |  |
| --- | --- |
| Oligonucleotides | Sequence (5'→3') |
| 3' Amine modified Poly(T) tailed sequence | 5'-TTTTTTTTTTTTTTTTTTTTTTTTTTTTTTTT-3'NH<sub>2</sub>C6T |
| U6-specific LNA probe | 5'-CTCGCTTCGmGCAGCmACA-3'-6-FAM |
| miR-16-5p specific LNA probe | 5'-GTmAGCAGmCACGTAAATA-3'-6-FAM |

### 2.2. Study design and sample collection

In this study, the samples were collected from the subjects. The blood samples were collected by venipuncture after obtaining written informed consent. The blood samples (n=9) collected from individuals were allowed to stand for 60 min, and plasma was separated to isolate circulating cell-free miRNA (ccf-miRNA).

### 2.3. Isolation and characterisation of ccf-miRNAs

Extraction of ccf-miRNA using Nucleospin miRNA Plasma Kit (Macherey Nagel, Germany) was executed in accordance with the manufacturer’s instructions and protocol. Briefly, denaturation of the plasma samples in the lysis buffer was followed by precipitation with protein precipitation buffer, and the samples were then centrifuged. After centrifugation, the supernatant was collected, mixed with isopropanol, and loaded onto the NucleoSpin miRNA column. Following centrifugation, the miRNA-bound column was treated with rDNase and incubated at room temperature for 15 min to digest DNA. The column was washed and dried. The miRNA was eluted using RNase-free water. The concentration and purity of miRNA were assessed using a NanoDrop, and the aliquots were stored at -80°C for further use.

### 2.4. Polyadenylation and characterisation of isolated miRNA

The isolated miRNAs were subjected to poly-adenylation reaction using the Invitrogen poly-A tailing kit (Thermofisher Scientific, USA). The poly-A-tailed ccf-miRNAs were then subjected to cDNA synthesis using the Reverse Transcription System (Promega, USA) according to the manufacturer’s instructions. Following cDNA preparation, the ccf miRNAs were amplified by using Taq 2X Master Mix (NEB, US) and primers (**F-**CCACTCTAGCAGCACGTAAAT, **R-** TCACACTAAAGCAGCACAGTAA). The data were analysed by gel electrophoresis based on band intensity. Furthermore, the miRNA expression patterns were characterised using real-time PCR. The cDNA was prepared using the reverse transcription system, followed by amplification of the target miRNA. Insta Q96 PCR platform (Himedia, India) was used for the real-time PCR reaction. Relative expression of circulating miR-16 was quantified using the comparative *Ct* (2^-ΔΔCt^) method, with U6 serving as the endogenous normalisation control. To satisfy the assumption of normality for parametric testing, statistical comparisons were executed strictly on the log-transformed ΔCt values.

### 2.5. Fabrication of the capture facet of the array

The nano-polystyrene comprises spherical structures with surface-bound carboxylic groups, making them electrostatically stable. For homogeneous attachment of the poly(T)-tailed oligonucleotide sequence to the surface of the nano-polystyrene, the carboxyl groups were pre-activated using the EDC-NHS protocol. Prior to activation, a suspension (4% w/v) comprising of 2.204 × 10^9^ billion nano-polystyrene spheres was subjected to washing with 1 mL washing buffer followed by centrifugation at 10,000 × g for 10 minutes. After removal of the supernatant, 50 μL of EDC solution and 50 μL of NHS solution, freshly prepared in activation buffer (50 mM MES), were added and vortexed. This mixture was incubated for 20 minutes at room temperature. To this solution, 20 μL of 3’end amine-modified poly(T) tailed oligonucleotide solution (10 µM) was added. Subsequently, a second aliquot of 50 μL each of EDC and NHS solutions was added to the above mixture to facilitate efficient binding. The final mixture was vortexed thoroughly and incubated for 3 hours at room temperature with intermittent vortexing. After 3 hours of incubation, a 0.005% v/v Tween-20 solution was added, and the resulting solution was centrifuged at 500 × g for 5 minutes to pellet the conjugated nano-polystyrene spheres. The supernatant was carefully removed, and the conjugated nano-polystyrene spheres were resuspended in 300 μL of storage buffer. The prepared conjugated nano-polystyrene spheres were stored at 4°C for further use **(Figure 1)**.

**Figure 1:**
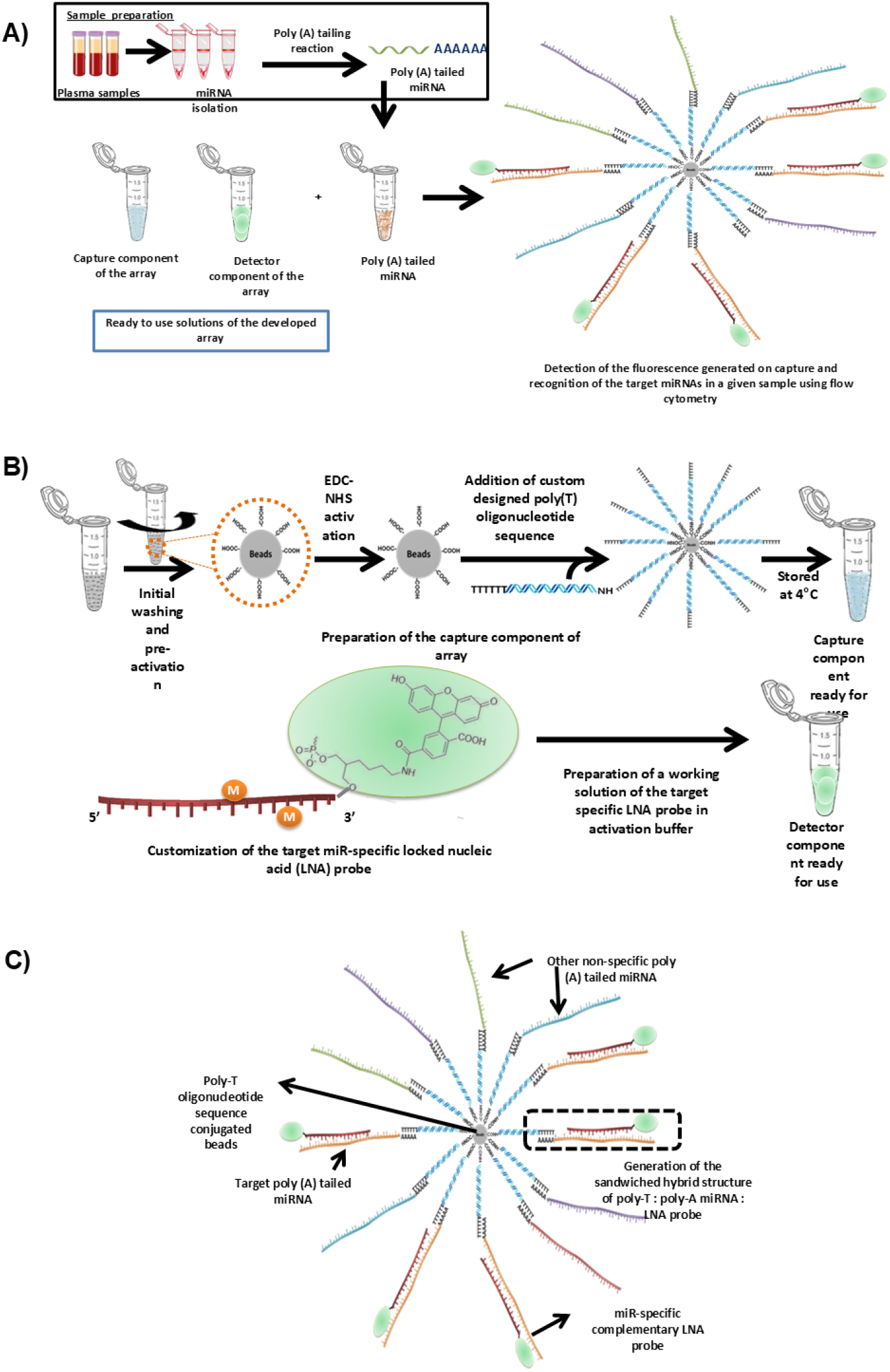
**A)** Schematic representation of the processes involved in the array methodology for the determination of the ccf-miRs in a given sample using flow cytometry. **B)** Schematic illustrating the hybridisation reactions involved in the preparation of a sandwiched nano-hybrid structure with respective facets for the determination of specific ccf-miRs. **C)** Schematic illustration of the hybridisation and detection of the target poly-A-tailed ccf-miRs by the developed nano-polystyrene-based method following complementary base pair matching.

### 2.6. Detection of target ccf-miRs

For the analysis of ccf-miRs of interest in a given sample, 20 μL of conjugated nano-polystyrene and poly-adenylated ccf-miRs sequences were allowed to incubate at 65 °C for 8 minutes. This step is crucial for the linearization of polyadenylated ccf-miRs and for efficient binding to the poly-T oligonucleotide sequences attached to nano-polystyrene. After incubation, the poly-A:poly-T hybrid-containing solution was immediately cooled on ice, washed with 500 μL of wash buffer, then centrifuged at 10,000 × g for 10 minutes, with the supernatant removed. 50 μL of 0.1X PBST and 20 μL of ccf-miR-specific probe solution (10 μM) were added after a 60-minute incubation at 50°C. The reaction was immediately cooled on ice. The obtained sandwich hybrid solution was then resuspended in 300 μL of 0.1X PBST for flow cytometry-based fluorescence measurement.

### 2.7. Applicability and selectivity of the developed array system

The detection capability of the prepared conjugates was analysed using flow cytometry in target-enriched samples and control samples. The enriched samples consisted of *ccf*-miRs isolated as per the manufacturer’s instructions. For the detection, 30 μL of each conjugate was added to 100 ng of previously isolated *ccf*-miRs in separate tubes. The isolated ccf-miRs were added to the tube of conjugates. The volume of each tube was adjusted to 300 μL with nuclease-free water (NFW), and the analysis was performed on a flow cytometer (Attune NXT, Thermo Fisher Scientific, USA). For selectivity, the conjugates were added to a solution containing different miRNAs, such as 29a, 7e, and 202.

### 2.8. Sensitivity of the developed array methodology

The sensitivity of a developed array is defined as the relationship between the change in analyte concentration and the transducer’s signal intensity. Ideally, a developed array should generate a signal in response to small fluctuations in the concentration of the target analyte. The developed methodology was evaluated for the feasibility of the array as a routine analytical approach for the sensitive estimation of minuscule quantities of target ccf-miRs in the intricate samples obtained after isolation from plasma.

### 2.9. Colocalization of developed array methodology with targets

To calculate colocalization, the developed array images were acquired using different channels along the same axis at equal intensities. Following image completion, the image was examined using the ImageJ programme. In the initial stage of ImageJ, two monochromatic images (A and B) were loaded into the platform. Yellow fluorescence revealed potential colocalization between the two images. A scatter plot was plotted, comprised of dots, appearing as a cloud, indicating complete colocalization.

### 2.10. Statistical Analysis

Each result’s mean and standard error were calculated. The GraphPad Prism Version 5.03 was used for statistical analysis. Differences in expression between the NSCLC, (n=10) and healthy control (n=5) cohorts were evaluated utilising an unpaired, two-tailed Student’s t-test. Data are expressed as the mean ± Standard Error of the Mean (SEM). Statistical significance was defined as *p*<0.05. Graphical visualisations and statistical computations were performed using Python (SciPy and Matplotlib).

## 3. Result and Discussion

miR-16-5p was selected as the target biomarker owing to its well-established role in the initiation and progression of NSCLC. Previous studies have demonstrated that dysregulation of miR-16-5p is associated with key oncogenic processes, including uncontrolled cellular proliferation, resistance to apoptosis, angiogenesis, and metastatic progression. Furthermore, altered expression levels of circulating miR-16-5p have been reported in plasma and serum samples obtained from NSCLC patients, indicating its potential utility as a non-invasive biomarker for early diagnosis, prognosis, and therapeutic monitoring (14, 15). Comparative expression analysis using RT-PCR and the 2^−ΔΔCt method following normalisation with the endogenous control U6, demonstrated a statistically significant difference in circulating miR-16-5p expression between the NSCLC and healthy control groups (p < 0.05), indicating that the expression of this miRNA is markedly altered during NSCLC development **(Figure 2)**. This differential expression is consistent with the reported regulatory role of miR-16-5p in multiple cellular processes, suggesting that its dysregulation reflects the underlying molecular alterations associated with NSCLC. The significant variation in circulating miR-16-5p expression also highlights its potential as a clinically relevant liquid biopsy biomarker. Since circulating miRNAs are released into the bloodstream and remain relatively stable, changes in their expression can provide valuable insights into disease status without the need for invasive tissue sampling. The observed expression profile of miR-16-5p therefore supports its potential utility in distinguishing NSCLC patients from healthy individuals. Unlike conventional tissue biopsies, ccf-miRNAs can be obtained via minimally invasive liquid biopsy, thereby offering a convenient approach for repeated disease assessment. Therefore, the development of rapid, sensitive, and amplification-free approaches for detecting circulating miR-16-5p is of considerable importance for improving NSCLC diagnostics (16, 17). To address these challenges comes with conventional methods, a nano-hybrid array was successfully developed for the selective detection of circulating miR-16-5p using poly(T)-functionalized nano-polystyrene particles and a 6-FAM-labelled fLNA probe. The platform facilitated efficient sandwich hybrid formation, enabling specific target capture and fluorescence-based detection. The high affinity of the fLNA probe and the large surface area of the nano-polystyrene particles contributed to enhanced hybridisation efficiency, allowing rapid and amplification-free detection of miR-16-5p. The incorporation of locked ribose conformations within the fLNA structure increases duplex stability and improves mismatch discrimination, enabling highly selective recognition of target miRNA sequences even in the presence of structurally similar nucleic acids. Upon hybridization of the fLNA probe with the capture d target miRNA, a sandwich nano-hybrid structure consisting of poly(T)-nano-polystyrene/poly(A)-tailed miRNA/LNA probe was formed. The formation of this ternary complex generated a fluorescence signal through the attached 6-FAM fluorophore, allowing direct quantification of target miRNA abundance.

**Figure 2.**
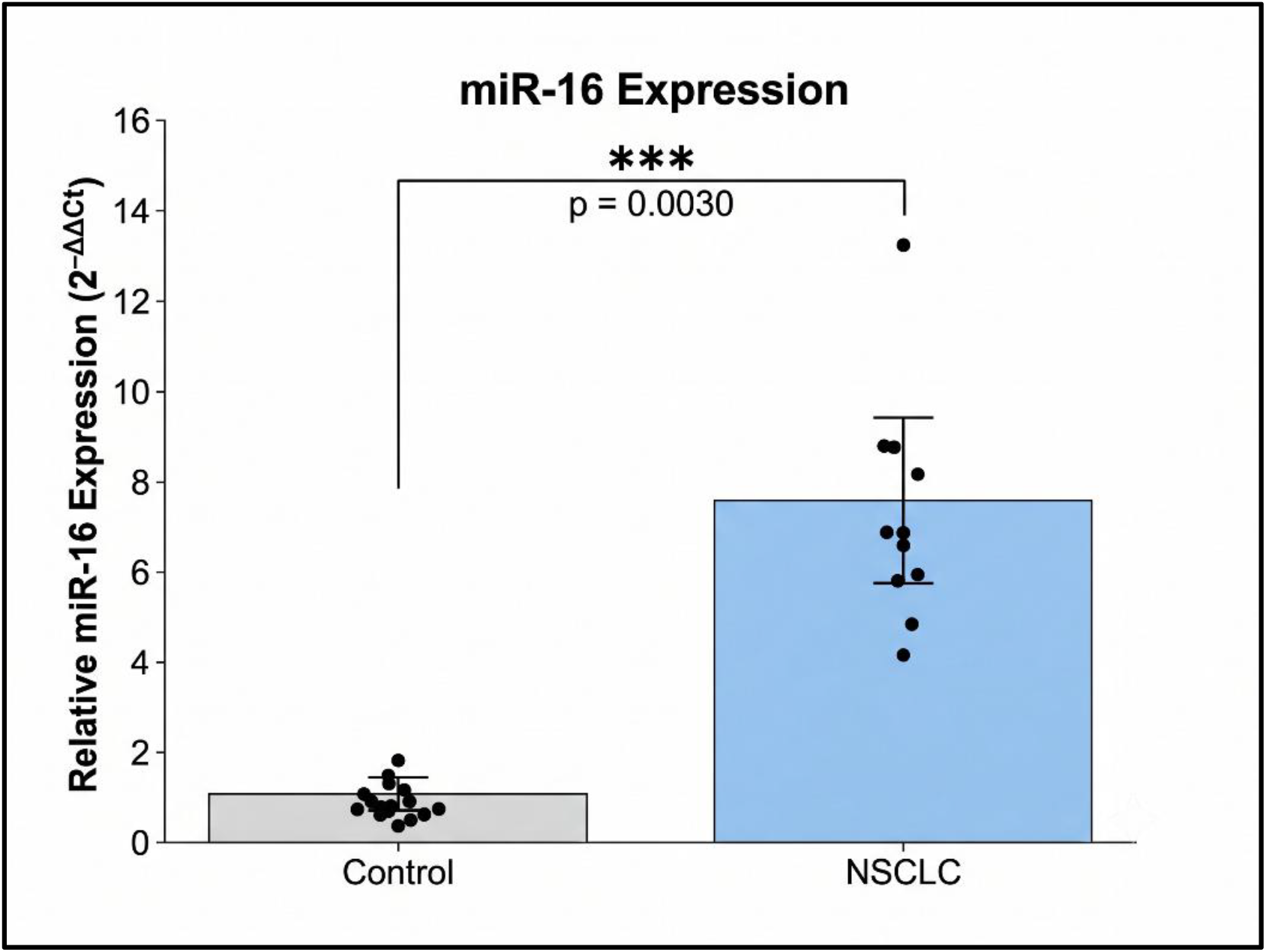
The Relative expression of circulating miR-16 in NSCLC. The fold change of miR-16 expression was evaluated in NSCLC samples (n=10) compared to healthy controls (n=5). Expression levels were normalised to the endogenous reference gene U6 using the 2-ΔΔCt method. Data are presented as mean ±SEM. Statistical significance was determined using an unpaired t-test; *** denotes p<0.01(p=0.0030).

The analytical performance of the developed nano-hybrid platform can be attributed to the complementary roles of the nano-polystyrene capture matrix and the fLNA recognition probes. The high surface area of the nano-polystyrene particles provides a greater density of immobilised capture oligonucleotides, thereby increasing the probability of target miRNA binding and improving hybridisation efficiency. In addition, the incorporation of fLNA probes enhances duplex stability and mismatch discrimination, allowing highly selective recognition of the target miRNA even in the presence of structurally similar nucleic acid sequences. These characteristics collectively contribute to efficient signal generation while minimising non-specific interactions. From an analytical perspective, the proposed sensing strategy demonstrates that combining nanomaterial-assisted target enrichment with high-affinity molecular recognition can overcome several limitations associated with conventional miRNA detection methods. By eliminating reverse transcription and nucleic acid amplification, the assay reduces sample manipulation and potential amplification bias while simplifying the overall analytical workflow. The performance of the developed array was assessed using flow cytometry, which included a blank, a negative control, and a positive control. The samples devoid of nucleic acids served as negative controls, while the samples enriched for miR-16-5p and U6 served as positive controls. All these samples were tested against a blank solution that contains no capture or detector facet (i.e., blank nano-polystyrene). This was necessary to analyse and nullify the auto-fluorescence (if any) generated by the conjugated nano-polystyrene to prevent the interference with the fluorescence intensity of the developed array. Initially, blank was analysed in the Attune NXT flow cytometer to adjust gating and voltage settings. For the analysis, the BL1-H channel was set to 488-nm excitation with the 530/30-nm bandpass filter. To assess the nano-hybrid performance, blank, negative control and positive control were analysed. The negative controls demonstrated no shift in the fluorescent intensity due to the absence of ccf-miRs available for binding. Upon addition of the positive control sample, a significantly observable shift in the fluorescence intensity was observed in the samples enriched in target ccf-miR species.

The optimal hybridization between poly(T)-functionalized nano-polystyrene and poly(A)-tailed miRNA-16-5p was achieved within 8 min at 65°C, followed by incubation with the complementary fLNA probe at 50°C for 60 min. Among the tested probe volumes, 50–100 μL of 6-FAM-labelled LNA probe produced the highest fluorescence response, indicating efficient formation of the sandwich hybrid complex. Similarly, sample volumes ranging from 1–20 μL yielded higher signal intensities than larger volumes, suggesting improved hybridisation efficiency under these conditions. These optimised parameters were subsequently employed for all analytical studies.

The selectivity of the assay was assessed by comparing fluorescence responses from target-enriched samples, other miRNAs (miR-29a, 7e, and 202c), and blank controls using fluorometry. Negligible fluorescence was observed in the blank nano-polystyrene conjugates, confirming minimal background interference. In contrast, other miRNAs, along with miR-16-5p-enriched samples, exhibited great fluorescence intensities. However, the miR-16-5p shows the highest fluorescence intensity. The substantial increase in fluorescence signal observed exclusively in the presence of the complementary target sequence miR-16-5p demonstrates the high specificity of the LNA probes and confirms the array’s selective recognition capability **(Figure 3)**.

**Figure 3.**
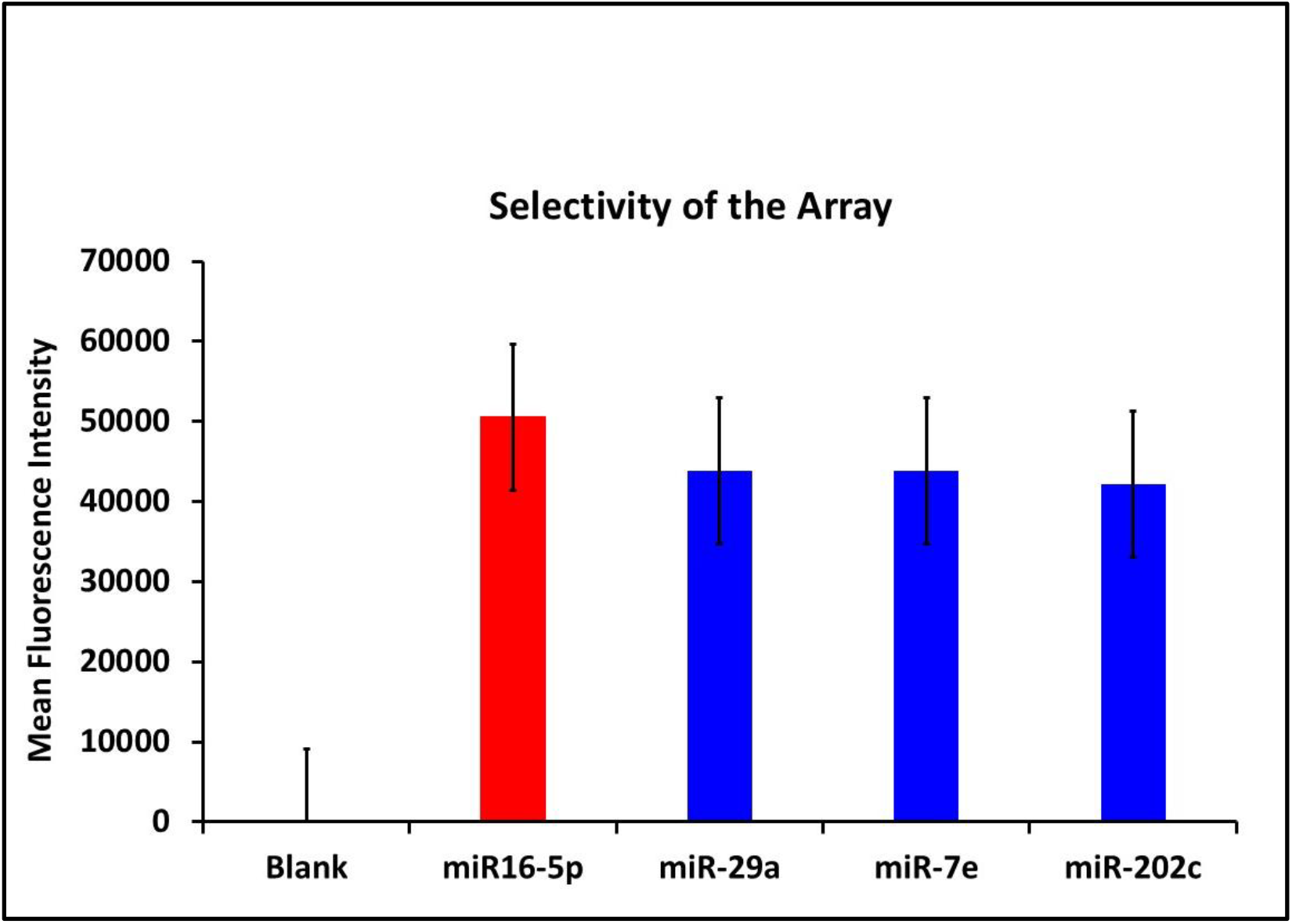
Graph showing the mean fluorescence intensity measurements (n=3) performed to evaluate the selectivity of the proposed methodology in the target ccf-miRs (miR-16-5P, miR-29a, miR-7e and miR-202c) enriched samples.

To evaluate the practical applicability of the platform, plasma-derived circulating cell-free miRNA samples were analysed following poly(A) tailing. The developed assay successfully identified target miRNAs even in the presence of numerous endogenous miRNA species. Samples containing 1 ng of isolated ccf-miRNA produced fluorescence intensities of 49.72% for U6 and 47.13% for miR-16-5p, while blank samples remained at baseline levels. These observations demonstrate that the nano-hybrid platform can selectively detect target miRNAs within complex biological matrices **(Figure 4)**.

**Figure 4:**
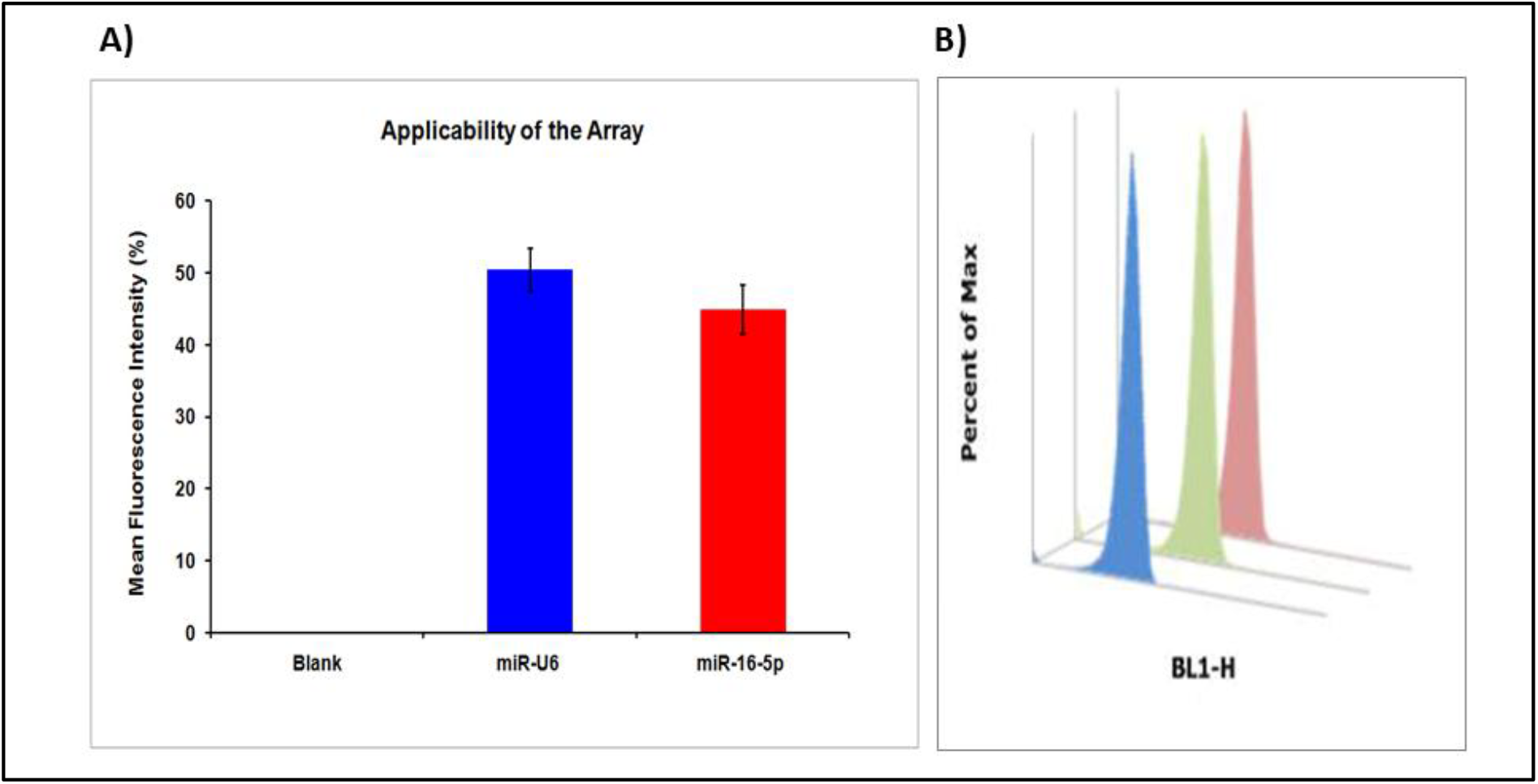
**A)** Graph representing the mean fluorescence intensity measurements (n=3) performed to evaluate the applicability of the proposed array method in detection of the target ccf-miRs (U6 & miR-16-5P) within a pool of ccf-miRs. **B)** Representative flow cytometry histograms showing the applicability of the array methodology in the determination of ccf-miRs of interest.

The sensitivity of the assay was investigated by decreasing the concentration of isolated plasma-derived miRNAs from 1 ng to 0.001 ng. Remarkably, the developed platform maintained strong fluorescence responses across the entire concentration range. For miR-16-5p, fluorescence intensities varied between 49.36% and 52.47%, whereas U6 detection yielded values ranging from 53.89% to 58.10%. Notably, even the lowest concentration tested (0.001 ng) produced measurable fluorescence signals well above background. These findings demonstrate the developed methodology’s ability to detect extremely low concentrations of circulating miRNAs directly from biological samples. The reproducibility of the assay was assessed by performing triplicate measurements at different miRNA concentrations. For U6 detection, fluorescence intensities ranged from 46.18% to 60.13%, while miR-16-5p exhibited fluorescence intensities between 43.93% and 49.92%. The low standard deviations and standard errors observed across all tested concentrations indicate excellent repeatability and measurement consistency. The reproducibility results confirm the robustness of the nano-hybrid platform and support its potential as a reliable analytical tool for routine quantification of circulating miRNAs **(Figure 5)**.

**Figure 5:**
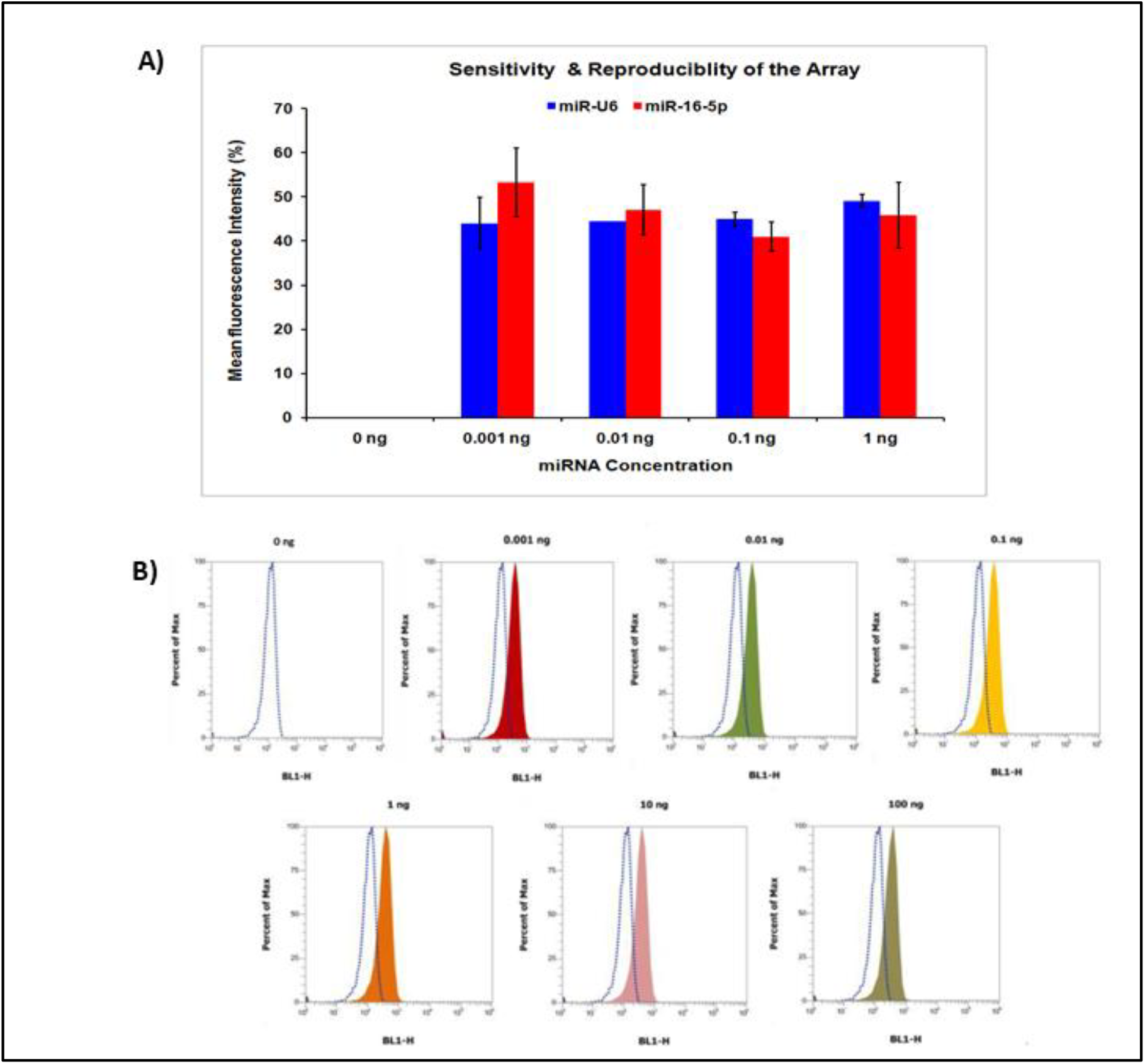
**A)** Graphical representation of the sensitivity and reproducibility of the proposed methodology. The mean fluorescence intensity values (n=3) were measured in different concentrations of target ccf-miR (U6 & miR-16-5P) in the given samples. **B)** Representative flow cytometry histograms showing the sensitivity and reproducibility tests of the array methodology in the determination of ccf-miRs of interest.

To evaluate the spatial association between the developed nanoanalytical framework and target miR-16-5p, coordinate-based co-localisation analysis was performed using high-resolution fluorescence microscopy. The nanoanalytical framework was visualised using differential interference contrast (DIC) imaging, while miR-16-5p-specific FAM-labelled probes and nucleic acid staining dyes were monitored through FITC and TRITC fluorescence channels, respectively [18-19]. Merged fluorescence images revealed prominent yellow regions, indicating substantial overlap between the fluorophore-labelled miR-16-5p and the nanoanalytical framework. The observed co-localisation confirmed successful target capture and hybridisation within the developed sensing platform. The high degree of co-localisation observed validates the efficient formation of the nanoanalytical complex and confirms the specificity of the developed detection strategy for the recognition of miR-16-5p. These findings provide direct microscopic evidence of successful target-probe interactions and support the robustness of the proposed platform for sensitive miRNA detection **(Figure 6)**.

**Figure 6.**
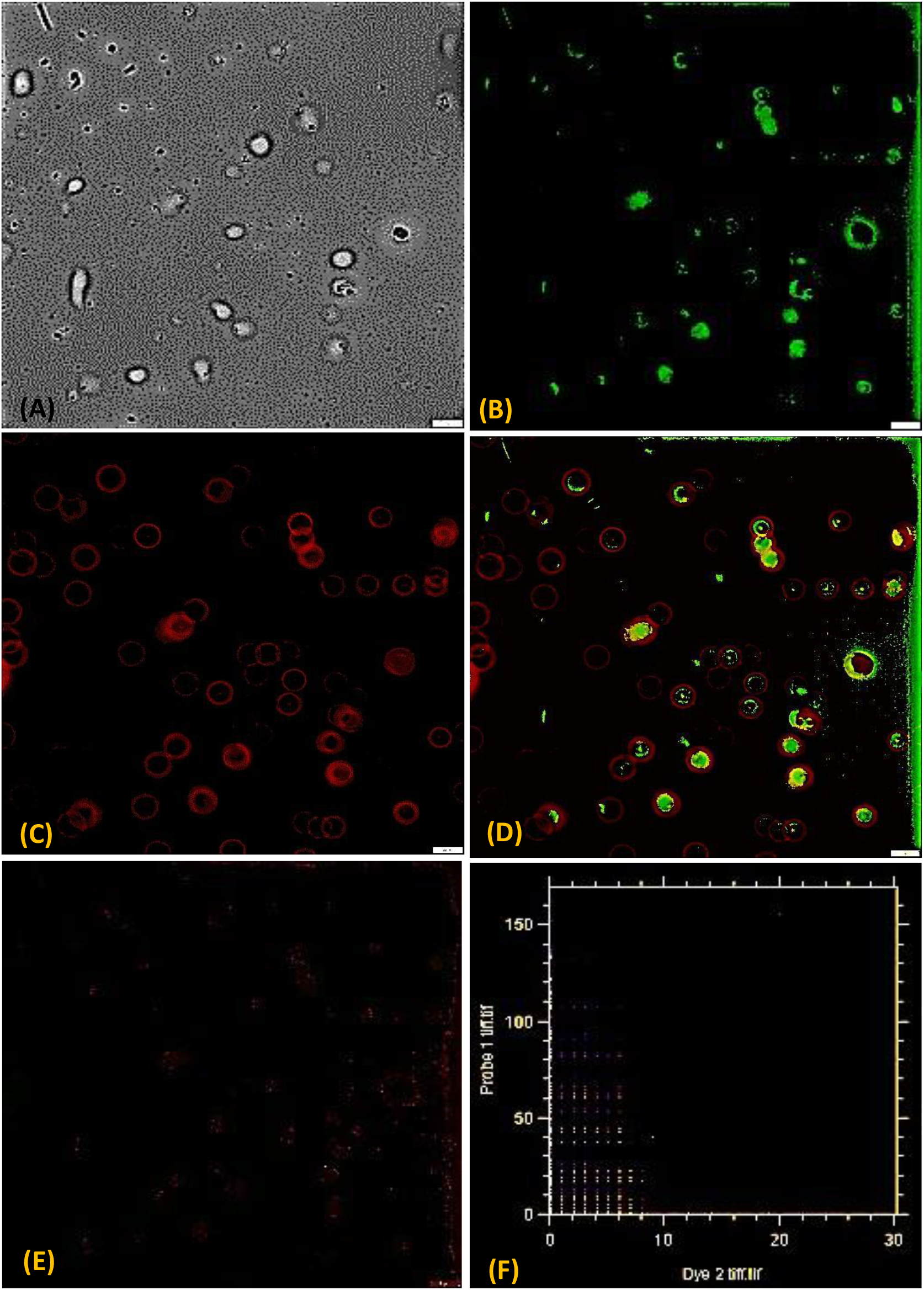
Nano-analytical framework, **(A)** Nano CML beads, **(B)** miR 16-5p labelled FAM conjugated with nano CML beads **(C)** DNA intercalating dye (Propidium iodide), **(D)** Composite image image depicting the overlapping between probe and intercalating dye, **(E)** Costes mask and **(F)** Scatter plot

## Conclusion

In the present study, a nano-hybrid array platform was successfully designed and assessed for the rapid and sensitive identification of NSCLC-associated ccf-miRNAs. The developed sensing strategy, based on poly(T)-functionalized nano-polystyrene particles and fluorescently labelled LNA probes, enabled efficient target capture and highly specific molecular recognition through sandwich hybrid formation. The assay demonstrated excellent analytical performance, including high selectivity, sensitivity, and reproducibility, for detecting miR-16-5p and U6 in plasma-derived circulating cell-free miRNA samples. All together, our findings indicated that the combination of nano-polystyrene-mediated target enrichment and LNA-based molecular recognition provides an effective approach for the direct detection of low-abundance ccf-miRNAs in plasma samples. Given the established association between ccf-miRNAs and NSCLC progression, the proposed methodology has significant potential for future applications in early diagnosis, disease monitoring, and assessment of therapeutic response [18]. The platform’s adaptability also promises its usage in designing multiplex detection assays for precision oncology and point-of-care diagnostic settings. Although such developments may certainly require extensive pre-clinical and clinical studies, before translation from bench to bedside.

## Acknowledgement

The authors thank the technical staff members, Mr. Pushpendra Gupta and Mr. Lalit Lodhi, for their assistance in collecting samples.

## Ethical Declaration

The study received approval from the Institutional Ethics Committee (IEC) of ICMR-NIREH under the reference number 5/13/1/PKM/ICRC/2020/NCD-III.

## Author Information

## Authors

**Ruchita Sandilya -** *Division of Environmental Biotechnology, Genetics & Molecular Biology (EBGMB), ICMR-National Institute for Research in Environmental Health (NIREH), Bhopal, India*

**Arpit Bhargava -** *Faculty of Science, Ram Krishna Dharmarth Foundation (RKDF) University, Bhopal, India*

**Pooja Ratre -** *Division of Environmental Biotechnology, Genetics & Molecular Biology (EBGMB), ICMR-National Institute for Research in Environmental Health (NIREH), Bhopal, India*

**Vikas Gurjar -** *Division of Environmental Biotechnology, Genetics & Molecular Biology (EBGMB), ICMR-National Institute for Research in Environmental Health (NIREH), Bhopal, India*

**Olga Goryacheva -** *Faculty of Materials Science, Shenzhen MSU-BIT University, Shenzhen 518172, People’s Republic of China*

**Rajnarayan Tiwari -** *Division of Environmental Biotechnology, Genetics & Molecular Biology (EBGMB), ICMR-National Institute for Research in Environmental Health (NIREH), Bhopal, India*

**Irina Yu Goryacheva -** *Institute of Chemistry, Saratov State University, Astrakhanskaya 83, 410012, Saratov, Russia*

## Authors Contribution: PKM devised the concept, developed the methodology, and supervised the experiments; RS, AB and PR performed most experiments; IYG characterized samples; and RS designed the figures; VG and PR performed the data analysis and interpretation; IYG and RT reviewed the manuscript; and PR, VG, RS, AB and PKM drafted the original manuscript

## Data Availability: The data supporting this study’s findings are available upon request from the corresponding author

## Abbreviations

ccf-miRNAs: Circulating cell-free microRNAs
NSCLC: Non-Small-Cell Lung Cancer
LNA: Locked Nucleic Acid
LC: Lung Cancer
LDCT: Low-Dose Computed Tomography
fLNA: Fluorophore-Labelled LNA
EDC: 1-Ethyl-3-(3-dimethylaminopropyl)carbodiimide
NHS: N-hydroxysuccinimide
SEM: Standard Error of the Mean
DIC: Differential Interference Contrast

